# The dual coding gene *SLC35A4* protects against oxidative stress

**DOI:** 10.1101/2024.12.14.627418

**Authors:** Ikram Ajala, Imadeddine Tiaiba, Benoît Grondin, Damien Lipuma, Souleimen Jmii, Hiba Benlyamani, Laurent Cappadocia, Benoît Vanderperre

## Abstract

Alternative proteins (AltProts) represent a newly recognized class of biologically active proteins encoded from alternative open reading frames (AltORFs) within already annotated genes. This study focuses on the *SLC35A4* gene, which encodes both the reference protein SLC35A4 and the alternative protein AltSLC35A4. Using a combination of microscopy and biochemical analyses, we confirmed the presence of AltSLC35A4 in the inner mitochondrial membrane, resolving previous conflicting reports. Previous studies employing ribosome profiling have revealed that during oxidative stress induced by sodium arsenite, the reference coding sequence of *SLC35A4* exhibits the largest increase in translational efficiency among all cellular mRNAs. Our results confirmed this translational upregulation, with the emergence of SLC35A4 protein isoforms during oxidative stress in an upstream ORF-dependent manner. Notably, the expression of AltSLC35A4 remained unchanged during oxidative stress. Knock- out of SLC35A4 enhanced sensitivity to oxidative stress in a rescuable manner, indicating a direct implication for SLC35A4 in stress resistance. In conclusion, our research provides compelling evidence for the functional significance of the dual-coding nature of *SLC35A4* for resistance to oxidative stress and highlights the importance of considering AltProts in the functional study of eukaryotic genes.

## Introduction

Over 21 years following the completion of the Human Genome Project, our comprehension of the "annotated" genome remains incomplete. The number of human protein-coding genes has remained largely unchanged since the publication by the Human Genome Sequencing Consortium ^1^. For decades, proteome exploration has been constrained by the belief that a eukaryotic gene generates a single, unique protein (splice isoforms excluded). However, numerous recent studies have disputed this notion, unveiling several classes of novel biologically active proteins involved in diverse physiological and pathological mechanisms^2–5^. These previously underestimated proteins are termed alternative proteins (AltProts) and are expressed in cells alongside the already identified proteins (referred to as reference proteins), thereby significantly expanding the known protein repertoire.

AltProts are encoded from alternative open reading frames (AltORFs) within previously annotated transcripts^6^ . They can be translated from RNAs previously considered non-coding, such as long non-coding RNAs (lncRNAs) and circular RNAs (circRNAs), from the untranslated regions (UTRs) of mRNAs, and may also overlap the coding sequences (CDS) of reference proteins but in a different reading frame ^2,6,7^. Consequently, an eukaryotic mRNA can function as polycistronic, encoding multiple proteins with entirely distinct primary sequences. Although the functions of several AltProts have been characterized, only a few research groups currently recognize their existence in their work. Given the relatively recent discovery of these proteins, they remain largely understudied ^2,6–8^, and the vast majority have yet to be functionally characterized.

An illustrative example is the double-coding gene *SLC35A4*. Located on chromosome 5q31.3, it encodes two proteins: AltSLC35A4, an AltProt encoded in the 5’UTR of the bicistronic mRNA, in addition to the reference protein SLC35A4 encoded by the reference CDS^6,9,10^. AltSLC35A4 is considered one of the first discovered alternative protein with its primary description and detection by mass spectrometry provided by Vanderperre et al^11^.

The SLC35 solute carrier family comprises members of the evolutionarily conserved human nucleotide sugar transporter (NST) family. The SLC35 solute carrier family of human NSTs is divided into 7 subfamilies (SLC35A-G), identified based on sequence similarity. Each NST subfamily is further subdivided to differentiate the type of substrate(s) presumed to be transported ^12^. Moreover, several members of the SLC35 subfamily have been designated as orphan transporters, as their substrate specificity and physiological functions remain to be determined^13^.

To date, the *SLC35A* subfamily has been extensively characterized. It is postulated that SLC35A1 is responsible for the import of CMP-sialic acid and CDP-ribitol into the Golgi apparatus^14,15^, SLC35A2 is essential for UDP-galactose import ^16^, while SLC35A3 is crucial for UDP-N-acetylglucosamine uptake ^17^. Ury et al. found that SLC35A4 imports CDP-ribitol in the Golgi, and is thus redundant with SLC35A1 ^15^. Ribitol plays a significant role in the glycosylation of alpha-dystroglycan (α-DG), a protein crucial in linking the intracellular cytoskeleton to the extracellular matrix and in muscle function^18–20^. Evidence of expression of SLC35A4 at the endogenous level is however limited: the OpenProt repository reports 5 unique peptides detected by MS, but recent attempts to detect this transporter by MS were unsuccessful^21^.

Cells react to stress through various mechanisms, which can include activating survival pathways or triggering programmed cell death to remove damaged cells^22^. Previous studies using ribosome profiling during oxidative stress (OS) and endoplasmic reticulum stress, induced respectively by sodium arsenite (SA) and tunicamycin, indicate that the translation of AltSLC35A4 is maintained albeit at lower levels than in absence of stress, and that the CDS of SLC35A4 undergoes the greatest increase in translational efficiency of all cellular mRNAs^23,24^. This characteristic is vital for proteins that play key roles in the integrated stress response^25^, whether they are protective or promote cell death, raising the possibility that *SLC35A4* encoded proteins regulate the stress response and cell susceptibility to stress.

In this study, we aimed at clarifying the basic features of AltSLC35A4, as well as studying the interplay between both proteins encoded in *SLC35A4*. We used microscopy and biochemistry approaches to confirm that AltSLC35A4 is a transmembrane protein of the inner mitochondrial membrane in human cultured cells. Using recombinant SLC35A4-expressing constructs, we show that the presence of the upstream ORF (uORF) is necessary for OS-induced expression of novel SLC35A4 short isoforms. Finally, we used CRISPR/Cas9-generated SLC35A4 knock- out cells, as well as corresponding rescued lines, to test the importance of SLC35A4 in the cellular susceptibility to OS. We demonstrate that SLC35A4 expression is necessary for protection against sodium arsenite-induced OS independent from the short isoforms. Collectively, our data establish AltSLC35A4 as an inner mitochondrial membrane integral protein and provide the first evidence of the functional importance of the *SLC35A4* gene in the response to OS.

## EXPERIMENTAL PROCEDURES

### Molecular cloning

All primers sequences are outlined in Supplementary Table 1. All PCR reactions were performed using Q5 DNA polymerase (New England Biolabs) according to the manufacturer’s protocol. PCR products were purified using the EZ-10 Spin Column DNA Gel Extraction Minipreps Kit (Bio Basic) and plasmids were purified using the EZ-500 Spin Column Plasmid DNA Maxi-preps Kit (Bio Basic). All constructs were sequence verified by Sanger sequencing (Genome Québec). Uniprot IDs of human *SLC35A4* encoded proteins are as follows: AltSLC35A4, L0R6Q1; SLC35A4, Q96G79.

For the AltSLC35A4-3xHA construct, a dsDNA fragment was synthesized (High-Q Strings, ThermoFisher Scientific) containing the whole 5’UTR of the *SLC35A4* mRNA including the AltSLC35A4 coding sequence fused to 3 C-terminal HA tags via a GGGGS linker, and flanked by regions homologous to the digested plasmid. The dsDNA fragment was inserted using Gibson assembly into the EcoRI and NheI restrictions sites of the pLJM1-EGFP backbone, replacing the EGFP coding sequence. To obtain the 3xHA-AltSLC35A4 plasmid, the 3xHA tags sequence (PCR 1), and the AltSLC35A4 coding sequence (PCR 2) were PCR-amplified from the AltSLC35A4-3xHA construct. In PCR 1, a start codon, C-terminal GGSG linker, and a 3’ terminal sequence overlapping the 5’ terminal sequence of PCR 2 were added. In PCR 2, a stop codon was added after the last coding codon of AltSLC35A4. Sequences homologous to the EcoRI/NheI digested pLJM1 plasmid were also added at the corresponding extremities. The amplicons were inserted in the plasmid in a single Gibson assembly reaction. To obtain the SLC35A4-3xHA construct, the SLC35A4 sequence was PCR-amplified (PCR 3) from a previously obtained plasmid containing the coding sequence of SLC35A4 amplified from the genomic DNA of 143B cells. A second amplicon containing a GGGGS linker followed by 3xHA tag and a stop codon was PCR-amplified from the pLJM1_hAltSLC35A4-3xHA plasmid (PCR 4). The two fragments were assembled using Gibson assembly, and the assembled construct was PCR amplified using the forward and reverse primers from PCR 3 and PCR 4, respectively. The assembled SLC35A4-3xHA sequence was then cloned into the pLJM1 plasmid using T4 ligase at the EcoRI/NheI restriction sites. To clone the bicistronic AltSLC35A4_SLC35A4-3xHA sequence into the pLJM1 plasmid, the *SLC35A4* cDNA sequence was PCR-amplified (PCR 5) from human cDNA (gift from Catherine Mounier’s laboratory). The products from PCR5 and PCR 4 (described above; GGGGS linker + 3xHA + stop codon) were assembled using Gibson assembly and the assembled construct was PCR amplified using the forward and reverse primers from PCR 5 and PCR 4, respectively. The final AltSLC35A4_SLC35A4-3xHA sequence was cloned into the pLJM1 plasmid using T4 ligase at the EcoRI/NheI restriction sites. For the AltSLC35A4 (untagged) construct, the AltSLC35A4 coding sequence was PCR amplified (PCR 6) from the AltSLC35A4-3xHA plasmid, and inserted into the EcoRI/NheI digested pLJM1 plasmid by Gibson assembly. For cloning the coding sequence of human AltSLC35A4’s N-terminus (amino acids 2-60) into pRSF-Trx (used for protein purification in *E. coli*), the corresponding sequence was PCR amplified (PCR 7) from a plasmid containing human AltSLC35A4 fused to EGFP (SLC35A4_pEGFP-N1^11^, gift from Xavier Roucou’s laboratory). The amplicon was inserted into a BamHI-digested homemade pRSF-Trx vector.

### Cell culture and sodium arsenite treatment (SA)

143B and HEK293T cells were cultured in Dulbecco’s Modified Eagle Medium (DMEM) (Wisent, #319-005-CL) containing 10% fetal bovine serum (FBS) (Wisent, #080-450) and 1% penicillin-streptomycin (Bio Basic, #PB0135) under standard conditions (37°C in a humidified atmosphere with 5% CO2). Cells were treated with sodium arsenite (SA; Sigma, #7784-46-5) by diluting a 4 mM stock solution (prepared with ddH2O) into complete DMEM for the indicated durations. When immunofluorescence was performed after OS, cells were treated with SA for 3 h to prevent cells detachment during the staining procedure. Mock-treated cells were treated with the same volume of ddH2O. Note that 143B cells stably expressed Blasticidin-selectable Cas9 nuclease (obtained using the lentiCas9-Blast plasmid), which allowed generation of knock-out lines using CRISPR/Cas9.

### Lentiviral productions

Six-well plate were seeded with 1 million HEK293T. The following day, cells were transfected using Calcium Phosphate precipitation. The transfection mixture contained 2 µg of expression plasmid, 1 µg of pMDL, 0.6 µg of pMD2g, and 0.5 µg of pRSV-Rev, along with specified volumes of buffered H2O (pH 7.3), 2.5M CaCl2, and 2X HeBS (pH 7.00). Fifteen minutes were allowed for precipitate formation before adding to the cell medium. After 8 hours, the medium was exchanged with complete DMEM supplemented with non-essential amino acids (NEAA, Wisent, # 321-011-EL). The first supernatant harvest was conducted the next morning, and fresh DMEM with NEAA was added to the cells. A second supernatant harvest was performed the following morning. Supernatants were pooled and filtered (0.45 µm) and used to transduce recipient cells or kept at -80°C for future use.

### Stable cell line production

4.2 x10^4^ 143B cells/well were seeded in 12-wells plates in medium containing 4 µg/mL polybrene (approximately 10% confluency) and varying virus quantities for a total culture volume of 1 mL/well. 24 h post-transduction, cells were selected with 1.25 μg/ml puromycin. Wells where complete selection resulted in 70-90% cell death compared to untransduced control were chosen to ensure that most surviving cells statistically received only one lentiviral copy. These were kept as stable cell lines.

### Immunofluorescence

143B cells were seeded in 24-wells plates containing coverslips. The following day, cells were fixed using a 4% paraformaldehyde (PFA) solution diluted in PBS for 15 min, then washed three times in PBS. Subsequently, cells were permeabilized with 0.15% Triton X-100 in PBS for 5 min, followed by two additional PBS washes. Cells were then incubated in a blocking solution (5% normal goat serum/PBS) for 30 min at room temperature. The coverslips were then incubated overnight at 4°C or for 3 h at room temperature with primary antibodies (as indicated in Supplementary Table 2) in 2.5% normal goat serum/PBS. After three washes in 2.5% normal goat serum/PBS, cells were incubated for 1 h at room temperature in the dark with fluorescent secondary antibodies (Supplementary Table 2). Following three washes in PBS, nuclei were stained with Hoechst (Invitrogen, #H3570; 1/10,000 in PBS) for 10 min at room temperature, before 2 further PBS washes and coverslips mounting on glass slides. Images acquisition was carried out using a Nikon A1 confocal unit (20X or 60X-oil objective) and NIS-Element software (Nikon). Where indicated, images were acquired with an EVOS M5000 epifluorescence microscope.

### Western blotting (WB)

Cells in 6-wells plate were initially rinsed with PBS, scraped in PBS and collected in centrifuge tubes for 3 min at 1000 rpm. The pellet was rinsed a second time with cold PBS and after another centrifugation, the pellet was resuspended in either 1X Laemmli buffer with 2.5% β-mercaptoethanol or RIPA buffer (25 mM Tris-HCl pH 7.4, 0.1% SDS, 1% sodium deoxycholate, 1% NP-40, 150 mM NaCl, 1 mM EDTA, 10 µM leupeptin, 0.3 µM aprotinin), and incubated on ice for 20 minutes before sonication with a probe (20% intensity for 20 seconds). If RIPA was used, 5X Laemmli buffer containing β-mercaptoethanol was then added to the protein samples to achieve a final 1X concentration. Samples in 1X Laemmli were boiled at 95°C for 10 minutes before SDS-PAGE and transferred at 100 V in 20% EtOH-containing transfer buffer for 1 hour onto a nitrocellulose membrane (Millipore Sigma; 0.45 μm). For AltSLC35A4, samples were transferred at 0,15A in 20% MeOH for 2 hours onto PVDF membrane (Bio-Rad 0.22 μm). After blocking (blocking solution: 5% milk in TBS-T [0.1% Tween]) for one hour (overnight for AltSLC35A4), membranes were incubated with primary antibodies at 4°C overnight in blocking solution (2 hours for AltSLC35A4 at room temperature). Following three 5 min washes with TBS-T, membranes were incubated with HRP-conjugated secondary antibodies for 1 h in blocking solution. After three additional 5 min washes with TBS-T, the membranes were exposed to Clarity Western ECL (Bio-Rad) when stronger signals were needed. Chemiluminescent signal was visualized using the Fusion Fx7 machine (MBI).^6^ Densitometric analysis was performed using ImageJ software (U.S. National Institutes of Health, Bethesda, MD, USA), and values were corrected for loading using actin signal as a reference.

### Isolation of mitochondria-rich fraction, alkali treatment and proteinase K protection assay

Isolation of mitochondria-rich fraction in mitochondria buffer (MB: 210 mM mannitol, 70 mM sucrose, 1 mM EDTA, 10 mM HEPES [pH 7.5]) was done as previously, except that cells were disrupted by 2x 15 passages in a 25g 3/8 needle fitted onto a 1 mL syringe. For alkali treatment, 100 µg of the mitochondrial suspension was centrifuged at 13,000 rpm for 10 min at 4°C. The resulting pellet was resuspended in 100 µL of ice-cold 0.1 M Na2CO3 (pH 11.5) to achieve a concentration of 1 µg/µL, followed by a 20 min incubation on ice. After transfer to a pre-chilled Polyallomer Microfuge tube (Beckman), the sample was centrifuged at 55,000 rpm for 40 min at 4°C using a TLA-55 rotor to separate soluble and membrane-associated proteins (supernatant) from membrane-inserted proteins (pellet). The supernatant was transferred to a new tube, and 4X Laemmli buffer containing 10% β-mercaptoethanol was added to reach a 1X final concentration. The pellet was resuspended in 1X Laemmli Buffer containing 2.5% βME to reach the same volume as the supernatant sample. Samples were analyzed by WB.

Proteinase K protection assay was conducted as described previously^26^. Briefly, Mitochondria were prepared at a final concentration of 0.5 mg/mL in MB, swelling buffer (10 mM HEPES-KOH [pH 7.4]), or MB with 0.2% (v/v) Triton X-100. Proteinase K (PK) (Sigma-Aldrich #39450-01-6) at 10 mg/mL in water was then mixed gently by pipetting up and down to reach a final concentration of 50 µg/mL, and the mixture was incubated on ice for 20 minutes.

Following this, phenylmethylsulfonyl fluoride (PMSF) was introduced to a final concentration of 2 mM to inhibit PK activity. Samples were analyzed by WB.

### Expression of the N-terminus of AltSLC35A4 in *E. coli* for anti-AltSLC35A4 antibody generation

The sequence coding for hAltSLC35A4_2-60 was cloned into a home-made pRSF- Thioredoxin(Trx) vector and transformed into *E. coli* strain BL21 DE3 CodonPlus RIL. Cells were grown in Super Broth medium at 37°C until reaching an A600 of 1.0. Protein expression was performed at 18°C for 24 h using 0.1 mM isopropyl-β-D-thiogalactopyranoside (IPTG). Cells were collected by centrifugation, suspended in lysis buffer containing 20 mM Tris-HCl, pH 8.0, 500 mM NaCl, 20 mM Imidazole pH 8.0, 5 mM β-mercaptoethanol and 100 µg/mL lysozyme. Suspended cells were lysed by freezing and sonication. After centrifugation at 40,000 g for 30 min, the supernatant was applied to a Ni-QIAGEN Ni-NTA resin. Proteins were eluted with an elution buffer containing 20 mM Tris-HCl pH 8.0, 500 mM NaCl, 500 mM Imidazole pH 8.0 and 5 mM β-mercaptoethanol. Fused proteins were cleaved by Tobacco Etch Virus Protease to remove the Trx tag, and then purified by gel filtration chromatography (Superdex 200 pg, GE Healthcare) in PBS buffer. The concentration of purified protein was determined using Bradford assay and used by the MediMabs company for rat polyclonal antibody generation using their standard protocol. The rat serum was used directly for WB without further purification.

### CRISPR-Cas9 editing

Briefly, 143B cells were transduced with a Blasticidin-selectable lentiviral cassette allowing stable expression the Cas9 nuclease (Addgene #52962). EasyEdit sgRNAs were purchased from GenScript and resuspended as per the manufacturer’s instructions. 143B-Cas9 cells were reverse-transfected with using Hiperfect reagent (Qiagen). Briefly, two sgRNAs (15.5 μL each from a 3 μM stock) were placed in a well of a 24-wells plate, then a mixture of 4.25 μL Hiperfect diluted in 137.5 μL serum-free DMEM was added to the sgRNAs, and incubated for 10 minutes at room temperature. 1 mL of a freshly trypsinized cell suspension (24,250 cells/mL) was added to the well. The plate was incubated overnight at 37°C with 5% CO₂. The next day, the medium was changed. On day 4, cells were trypsinized and resuspended. One-quarter of each well was collected for gDNA extraction (QuickExtract, Lucigen Corporation, # QE09050) and editing assessment by PCR, and a portion was used for sorting individual cells into 96-wells plates to derive monoclonal knock-out (KO) lines. Initial KO verification in isolated monoclonal populations was performed by PCR (Supplementary Fig. 1A,D). For SLC35A4, the PCR product was blunt-cloned using T4 ligase into the pBluescript KS (-) plasmid at the EcoRV restriction site and sequenced using Sanger sequencing (T7 promoter primer). For AltSLC35A4, the clones’ allelic sequences were analyzed using MiSeq at the CERMO-FC genomics platform. Sequences of sgRNAs and primers used for KO validations are shown in Supplementary Table 1. The consequences of mutations in each allele are shown in Supplementary Fig. 1B for the AltSLC35A4-KO clone, and in Supplementary Fig. 1E for SLC35A4-KO clones.

### Measurement of cell proliferation

An equal number of cells (0.05 x 10^6) were seeded in 12-wells plates with complete DMEM and incubated at 37°C for 72 hours. During this period, one plate was taken every 24 hours, and cells were trypsinized and counted using an automated cell counter (LUNA-II, Logos Biosystems) according to the manufacturer’s instructions. The number of cells at time 0 h was set as 1, and the number at each subsequent time point (24 h, 48 h, 72 h) was normalized to the 0 h count.

### Measurement of cell viability

The CellTiter-Glo® 2.0 Assay (Promega, #G9242) provides a measure of cell viability in culture by quantifying ATP, which indicates the presence of metabolically active cells. The luminescence reading is directly proportional to the number of viable cells in culture. Briefly, an 3x10^4^ cells per well were plated in complete DMEM in a 96-wells plate compatible with the assay. The next day, the culture medium of each well group was replaced with 100 µL DMEM containing 40 μM SA (or vehicle for the control 0h group) for 0 h, 1 h, 2 h, 4 h or 8 h. At the end of the treatment, 100 μL of CellTiter-Glo 2.0 solution was added to all cell groups according to the manufacturer’s instructions, and the plate was analyzed using a luminescence reader. For each genotype, viability at each time point is expressed as percent viability relative to that at 0 h (defined as 100%).

### Bioinformatics and statistical analyses

DNA sequences of AltSLC35A4 from multiple species were retrieved from GenBank (http://www.ncbi.nlm.nih.gov/genbank) and aligned with Clustal Omega software (http://www.clustal.org). The results were subsequently visualized using Jalview. Re-analysis of mitochondrial proteomics data from mouse tissues^27^ was performed as previously^28^. Statistical analyses were performed using GraphPad PRISM.

## RESULTS

### *SLC35A4* is a dual-coding gene that encodes an alternative protein from the 5’UTR

In an early effort to identify alternative proteins encoded in the transcriptome of human and other eukaryotes using mass spectrometry, we previously detected AltSLC35A4 as an AltProt encoded within the 5’UTR of the *SLC35A4* mRNA^6^. The 103 codons of AltSLC35A4’s coding sequence spreads across 3 exons of the gene, whereas the reference protein SLC35A4 is exclusively encoded by exon 3 (Fig. 1A). Using tblastn analysis, orthologs of human AltSLC35A4 coding sequence were found in myelinated vertebrates, but not earlier in evolution. Conservation analysis was performed using ClustalO and visualized using the Jalview software. Our results show that AltSLC35A4 is highly conserved (66.02% identity between zebrafish (*Danio rerio*) and human; Fig. 1B). This suggests an important biological function in these organisms.

**Figure 1 :**
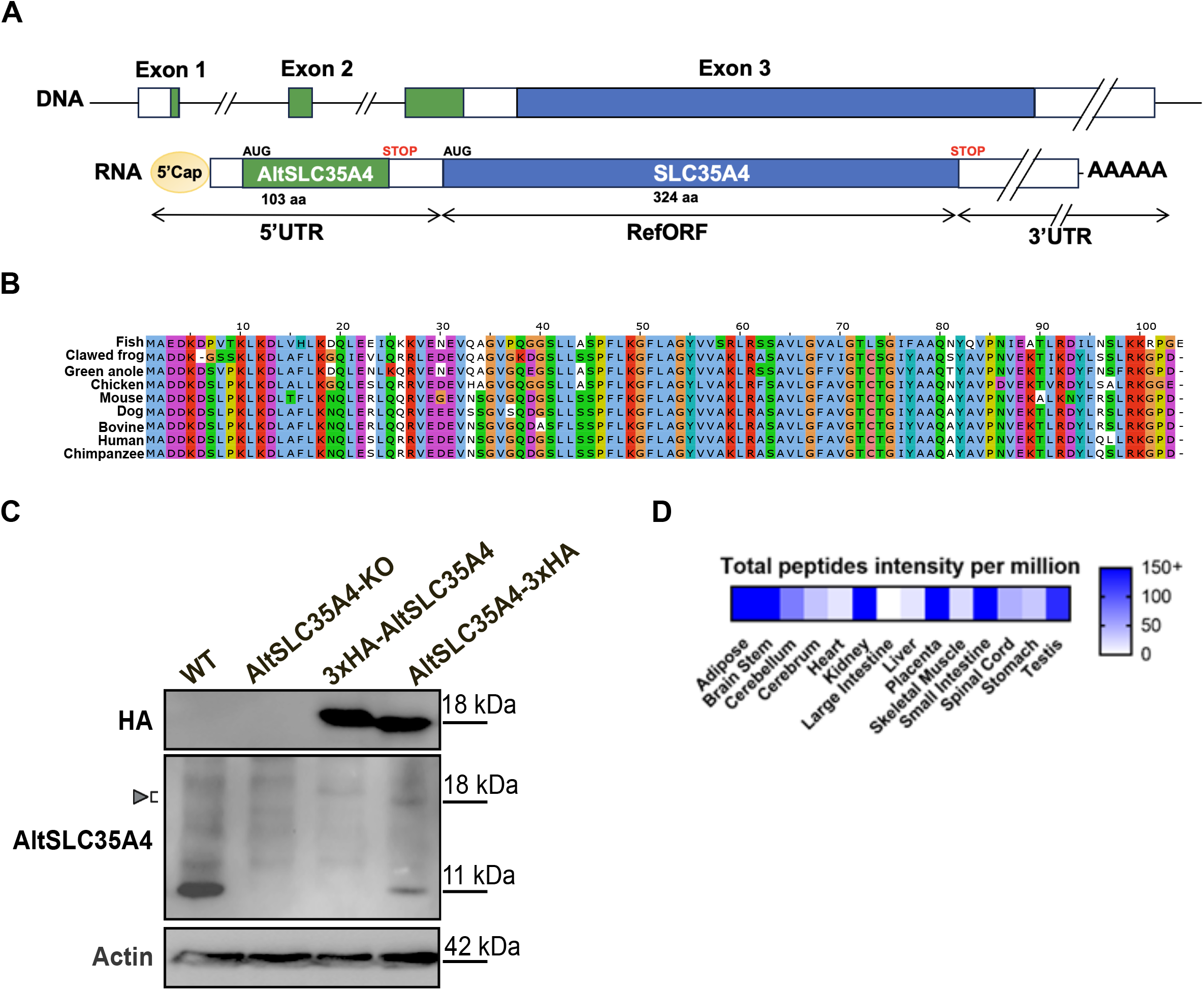
AltSLC35A4 is a highly conserved protein expressed from the *SLC35A4* bicistronic gene. **(A)** The *SLC35A4* gene is bicistronic, containing the canonical SLC35A4 CDS (in blue, +1 frame) and AltSLC35A4 CDS (in green, +2 frame). **(B)** Alignment of AltSLC35A4 protein sequences in zebrafish (*Danio rerio*), clawed frog (*Xenopus laevis),* green anole (*Anolis carolinensis),* chicken (*Gallus gallus*), mouse (*Mus musculus*), dog (*Canis lupus familiaris*), bovine (*Bos taurus*), human (*Homo sapiens*) and chimpanzee (*Pan troglodytes*). Residues are colored based on Jalview’s *Clustal* built-in colour scheme. **(C)** Detection of endogenous and exogenous expression of AltSLC35A4 with a home-made antibody in WT 143B cells, AltSLC35A4-KO, AltSLC35A4-KO stably expressing 3xHA-AltSLC35A4, and WT cells stably expressing AltSLC35A4-3xHA. **(D)** Reanalysis of mitochondrial proteomics data from 14 mouse tissues reveals abundant AltSLC35A4 expression in several organs^27^ .

### AltSLC35A4 is endogenously expressed in cell lines and tissues

We next sought to confirm that AltSLC35A4 is endogenously expressed using WB. In absence of a commercial antibody, we purified a recombinantly-expressed version of the N-terminal extremity of human AltSLC35A4 (amino acids 2-60) from *E. coli*, and used it to generate a custom polyclonal rat antibody. To ensure that the generated antibody accurately detects our protein of interest, we employed CRISPR/Cas9 genome editing to knock out (KO) AltSLC35A4 in human osteosarcoma 143B cells (AltSLC35A4-KO line, Fig. 1C). Our WB analysis showed that, in wild-type (WT) cells, the custom antibody recognizes a protein of approximately 11 kDa absent from KO cells, confirming the endogenous expression of AltSLC35A4 and specificity of our antibody (Fig. 1C). In parallel, in the KO line background, we generated rescued cells expressing AltSLC35A4 fused with 3xHA at the N-terminal extremity (3xHA-AltSLC35A4; also see Supplementary Fig. 1C). We also established a stable cell line that expresses AltSLC35A4 fused to a C-terminal 3x-HA tag (AltSLC35A4-3xHA) in a WT background. Both tagged versions were detected (albeit at very reduced levels compared to endogenous AltSLC35A4) at approximately 18 kDa, as expected (Fig. 1C), further confirming the specificity of the antibody.

Previous reports indicate that human AltSLC35A4 is localized in mitochondria^21,29^. To confirm expression of endogenous AltSLC35A4 in another vertebrate species and *in vivo*, and explore its expression pattern across several tissues, we reanalyzed a dataset containing the mitochondrial proteomics data from 14 different mouse tissues^27^, as performed previously^28^. AltSLC35A4 was detected in all tissues analyzed except the large intestine, with prominent expression in adipose tissue, placenta, kidney, and small intestine (Fig. 1D). These findings suggest that AltSLC35A4 is a conserved protein with significant expression in multiple tissues, indicating its potential importance in various biological functions.

### AltSLC35A4 is inserted in the inner mitochondrial membrane

We then sought to confirm the precise localization of AltSLC35A4 within mitochondria, to guide future functional studies regarding its molecular function. To investigate this, we used immunofluorescence and confocal microscopy in AltSLC35A4-3xHA and 3xHA- AltSLC35A4 expressing lines and confirmed colocalization with the mitochondrial marker TOM20 (Fig. 2A). AltSLC35A4’s localization in mitochondria was confirmed using subcellular fractionation, with both endogenous AltSLC35A4 and AltSLC35A4-3xHA enriched in the mitochondrial fraction (Fig. 2B). Rocha et al. used *in silico* analysis to predict that AltSLC35A4 contains a transmembrane domain between amino acids 62 and 84 ^21^. To biochemically confirm this, we subjected the mitochondrial fraction from AltSLC35A4-3xHA expressing cells to alkali treatment followed by ultracentrifugation to separate soluble and membrane-associated proteins (supernatant) from integral membrane proteins (pellet). Our results indicate that AltSLC35A4 is in the membrane fraction (Fig. 2C). We then sought to determine whether AltSLC5A4 is localized in the outer or the inner mitochondrial membrane. To achieve this, we conducted a proteinase K protection assay on purified mitochondria (Fig. 2D). In mitochondrial buffer, mitochondria are intact and proteinase K only degrades proteins exposed on the surface or the outer membrane (such as TOM20). However, AltSLC35A4 was degraded in a swelling buffer which allows access to the intermembrane space (where Smac/Diablo localizes) and inner membrane, or in fully disrupted mitochondria (Triton X-100 condition that sensitizes proteins from all compartments including matrix-localized SDHA). Our results clearly establish that AltSLC35A4 is inserted in the inner mitochondrial membrane.

**Figure 2 :**
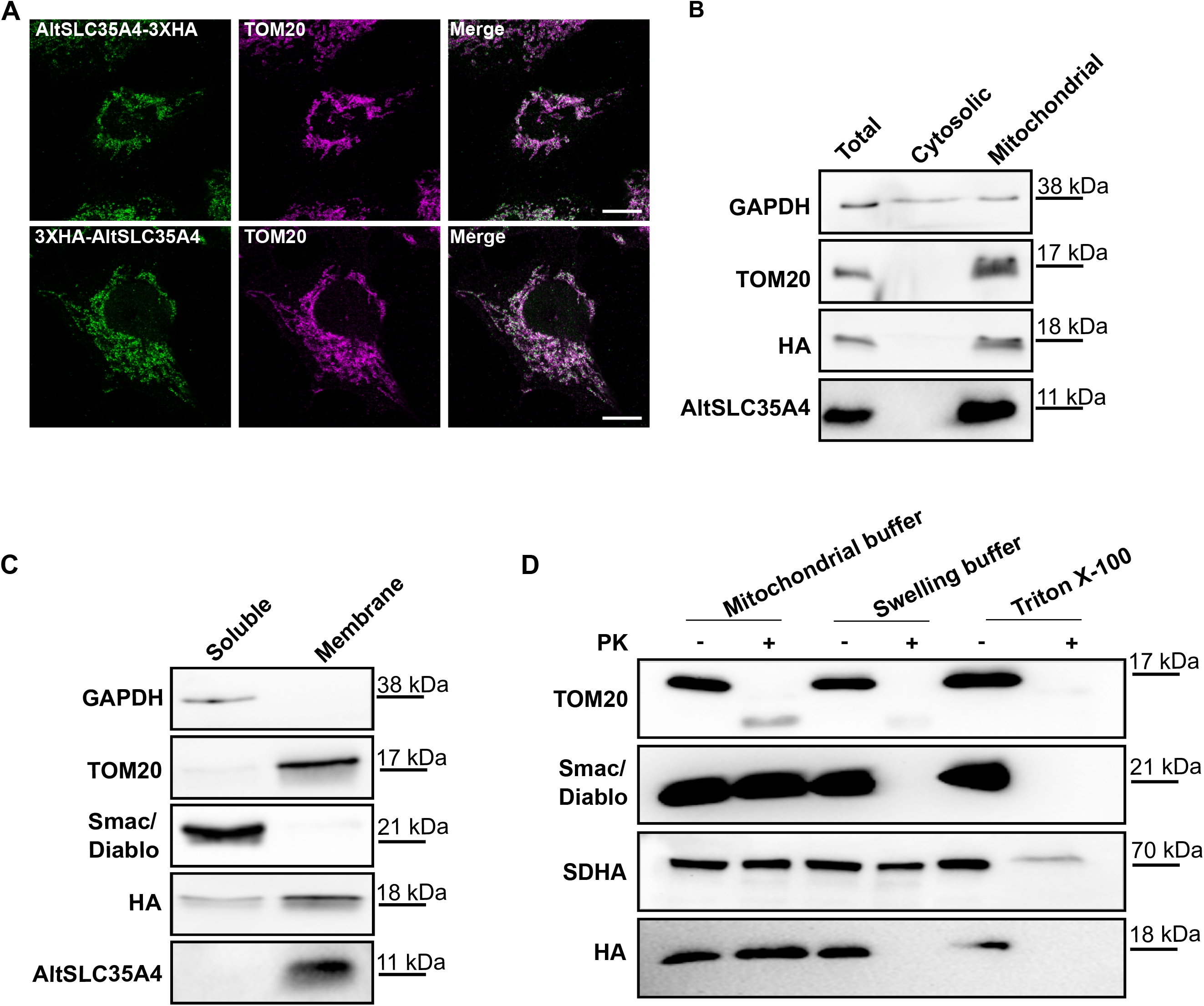
AltSLC35A4 is inserted in the inner mitochondrial membrane. **(A)** Immunofluorescence and confocal imaging on 143B cells stably expressing AltSLC35A4- 3xHA or 3xHA-AltSLC35A4 (green) shows that AltSLC35A4 colocalizes with the mitochondrial marker Tom20 (magenta). Scale bar: 10µm. **(B)** Cellular fractionation in AltSLC35A4-3xHA expressing cells shows that endogenous AltSLC35A4 and AltSLC35A4- 3xHA are enriched in the mitochondrial fraction. **(C)** Alkali treatment of the mitochondrial fraction from (B) followed by ultracentrifugation shows that AltSLC35A4 is an integral membrane protein. **(D)** Proteinase K protection assay on the mitochondrial fraction from 3xHA-AltSLC35A4 expressing cells reveals its localization in the inner mitochondrial membrane. The presented data is representative of N=3 independent experiments.

### Short isoforms are expressed in addition to full-length SLC35A4 in response to oxidative Stress (OS)

Having characterized AltSLC35A4, we then decided to deepen our understanding of the biological function of the *SLC35A4* gene, in particular in the stress response. Previous reports using ribosome profiling indicate that the coding sequence of the RefProt SLC35A4 undergoes the most important translational increase of all cellular mRNAs in response to OS and reticulum endoplasmic stress induced by sodium arsenite and tunicamycin, respectively^23,24^. Concomitantly, AltSLC35A4 translational efficiency was slightly reduced. We sought to test whether these changes in translational efficiency are reflected at the protein level. We were unsuccessful in detecting endogenous SLC35A4 by WB and immunofluorescence regardless of antibodies used, or by LC-MS/MS. To validate *SLC35A4’s* translational modulation at the protein level, we thus stably expressed a construct with a structure and sequence mimicking the endogenous *SLC35A4* mRNA, termed AltSLC35A4_SLC35A4-3xHA. In this construct, the cDNA of the dual coding *SLC35A4* gene is mimicked all the way to the stop codon of SLC35A4 (Fig. 3A). We excluded the 3’UTR region since translational regulation by uORFs is thought to be the most important contributor to translational upregulation in response to stress^23,30^. The combination of a strong promoter (CMV) with 3xHA C-terminal tagging of SLC35A4 allowed the detection of SLC35A4 expression by WB and immunofluorescence using an anti-HA antibody (Fig. 3A, 4A).

**Figure 3 :**
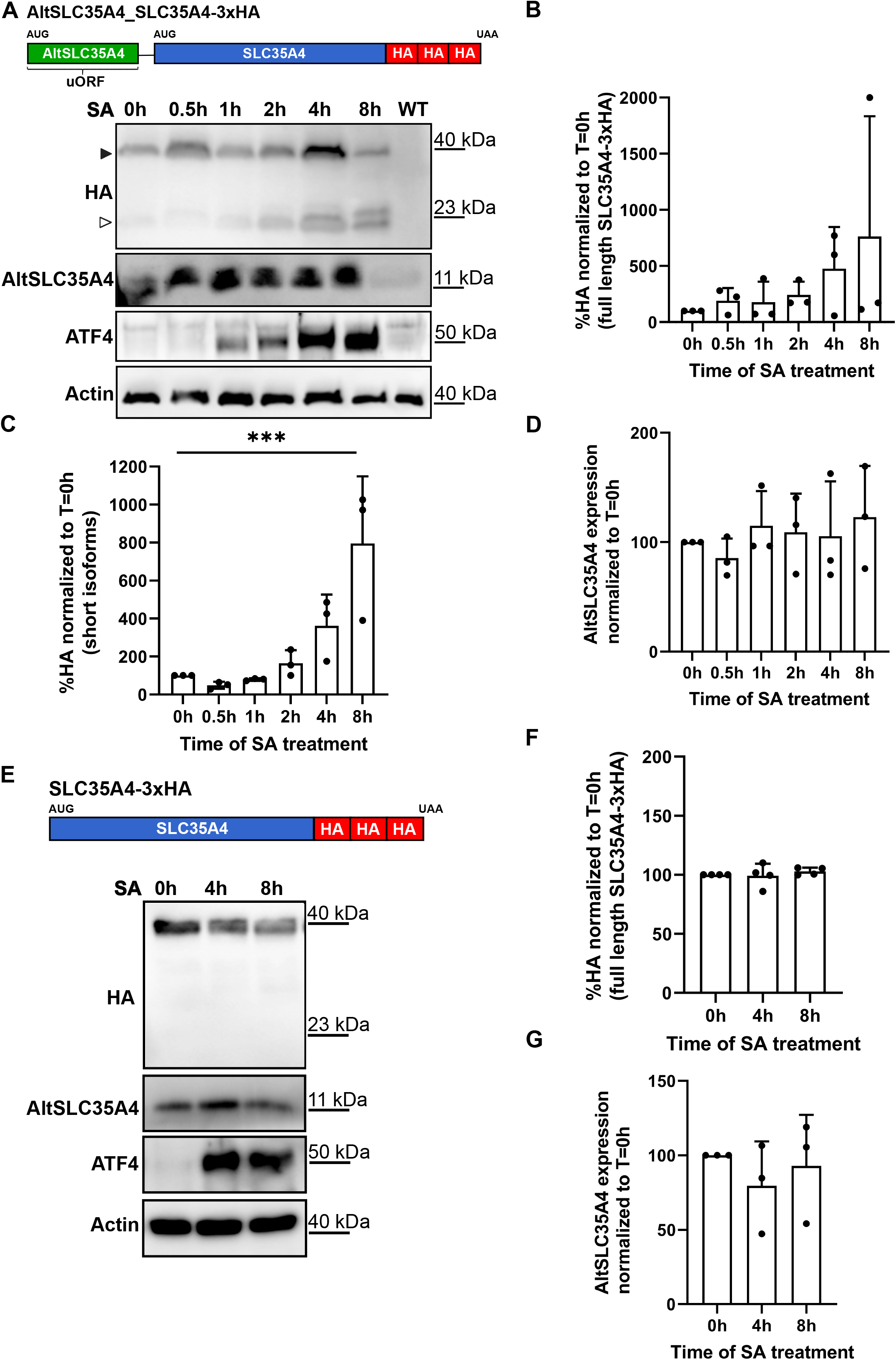
Translational regulation of SLC35A4 in response to oxidative stress in an upstream ORF-dependent manner. **(A)** Western blot analysis of 143B cells stably expressing the bicistronic AltSLC35A4_SLC35A4-3xHA construct and subjected to increasing times of 40 µM sodium arsenite treatment (0 h: vehicle). WT cells were also analyzed to ensure specificity of the HA signal. The full arrowhead indicates the full-length SLC35A4-3xHA protein, and the empty arrowhead indicates the shorter isoforms appearing in response to stress. SA: sodium arsenite. **(B- D)** Quantification of the Western blot analysis in (A), with expression levels normalized to actin and showed relative to the T=0h condition. **(E)** Western blots analysis of 143B cells stably expressing the SLC35A4-3xHA construct and subjected to increasing times of 40 µM SA treatment. **(F-G)** Quantification of the Western blots from (E). N=3-4 independent experiments. Statistical test: one-way ANOVA; *** P<0.001.

**Figure 4 :**
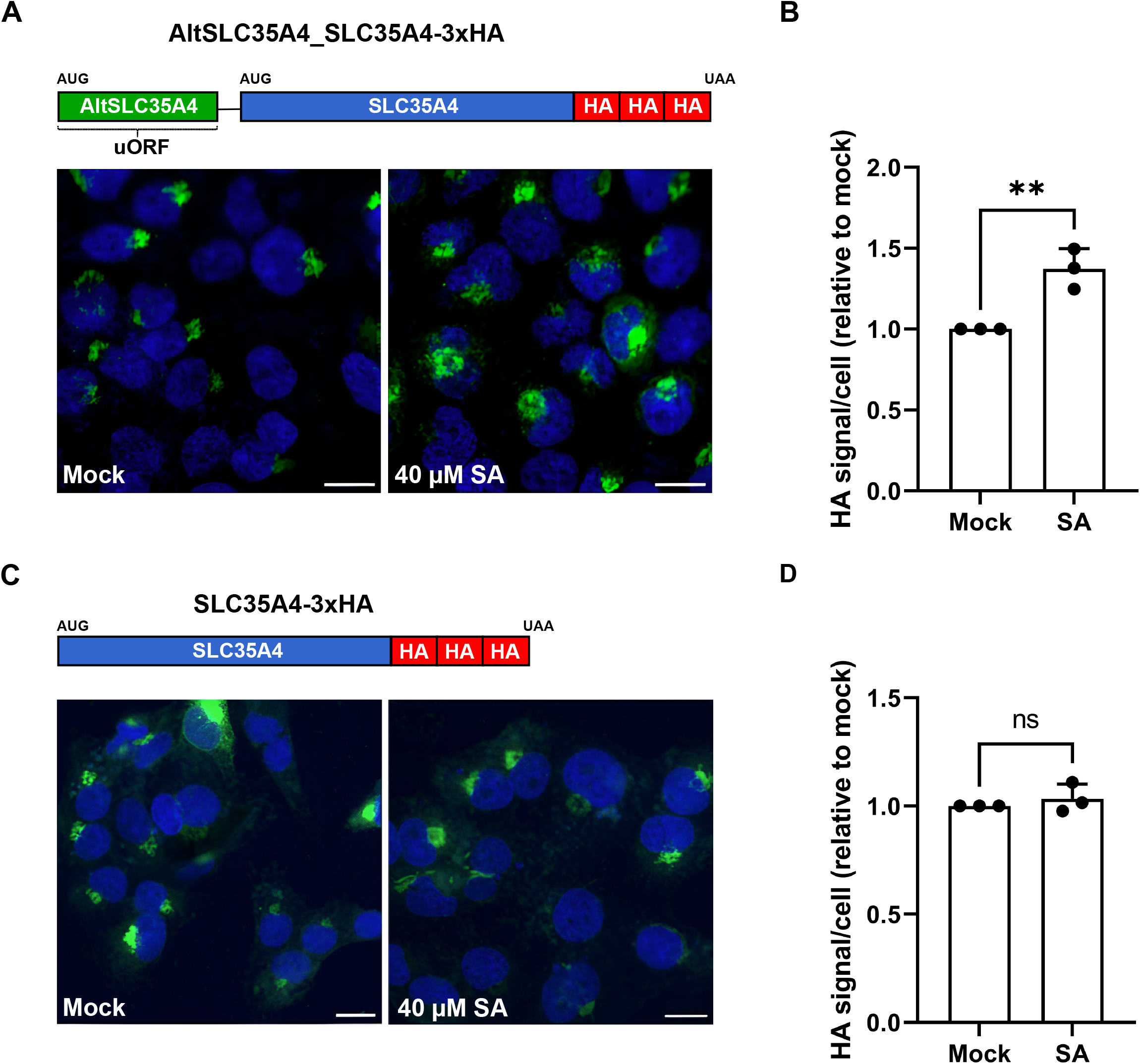
Total SLC35A4 increases in response to oxidative stress in a uORF-dependent manner. **(A,C)** Immunofluorescence and confocal imaging of 143B cells stably expressing the bicistronic AltSLC35A4_SLC35A4-3xHA construct **(A)** or the monocistronic SLC35A4- 3xHA construct **(C)**, treated or not for 3 hours with 40 µM sodium arsenite. The SLC35A4- 3xHA protein was labeled with an anti-HA antibody (green). Nuclei were stained with Hoechst (blue). Scale bars: 15 µm. **(B,D)** The HA signal intensity per cell from (A) and (C), respectively, was quantified with ImageJ and normalized to the HA signal of mock-treated cells. Statistical tests: unpaired t-test; N=3 independent experiments, n=24-44 cells; ** P<0.01.

To test if SLC35A4 protein levels are increased during OS, cells stably expressing AltSLC35A4_SLC35A4-3xHA were treated with sodium arsenite for various durations. WB analysis showed an increase in ATF4 expression with increasing treatment times, confirming the induction of OS (Fig. 3A). However, there was no significant difference in full length SLC35A4-3xHA levels between the different time points (Fig. 3A, full arrowhead, and Fig. 3B). Surprisingly, an increase in HA-containing bands of lower molecular weight was observed with increasing time of OS (Fig. 3A, empty arrowhead), reaching statistical significance after 8 h (Fig. 3B). The absence of HA signals signal in WT cells confirmed that HA-detected species were derived from the AltSLC35A4_SLC35A4-3xHA construct. To test the dependency of this phenomenon on the presence of an uORF, we performed a similar analysis in cells expressing a construct encoding solely SLC35A4-3xHA, without its endogenous upstream region (Fig. 3E, Supplementary Fig. 2B). In absence of the uORF, full-length SLC35A4-3xHA levels were also unchanged after OS, lower molecular weight bands were absent at baseline, and barely detectable after 8 h of OS (Fig. 3E,F). Our WB analyses were corroborated by immunofluorescence experiments. After 3 h of SA treatment, ATF4 became detectable in the cells nuclei, validating OS induction. The total SLC35A4-3xHA signal intensity per cell was increased upon OS when SLC35A4 was expressed from the uORF-containing construct AltSLC35A4_SLC35A4-3xHA (Fig. 4A,B), compatible with the appearance of additional isoforms observed by WB (Fig. 3A), but not from the monocistronic SLC35A4-3xHA construct (Fig. 4C,D). Finally, AltSLC35A4 levels did not vary in response to SA-induced OS, whether it was expressed from the endogenous locus (Fig. 3E,G), or from both the endogenous locus and the AltSLC35A4_SLC35A4-3xHA bicistronic construct (Fig. 3A,D). Overall, our data suggest that, while AltSLC35A4 levels appear maintained during OS, shorter isoforms of the SLC35A4 protein are produced in response to OS in a uORF-dependent manner, likely explaining the translational increase observed by ribosome profiling in previous studies^23,24^ and reducing the likelihood of proteolysis being the source of these additional isoforms.

### The reference protein SLC35A4 is important for cell viability in response to OS

Since translational induction despite general translation inhibition during stress is a feature of important stress response genes^22,25^, we then tested if the SLC35A4 reference protein is important for cellular resistance to stress. We generated two monoclonal SLC35A4-KO cell lines (cl. A, cl. B,) as well as their rescued counterparts expressing SLC35A4-3xHA from our monocistronic contruct (Supplementary Fig. 1D-F). Despite the absence of premature stop codons in mutated alleles, polypeptides produced from these alleles lack transmembrane domains 2, 3 and 4 (out of 10) and 8/15 residues predicted as important in forming the CDP-ribitol binding site (Supplementary Fig.3) ^15^. They were thus deemed dysfunctional. First, we confirmed that cell proliferation was unaffected in SLC35A4-KO lines in unstressed conditions using cell counting. Similar results were obtained for the AltSLC35A4-KO line (Fig. 5A). These results indicate that SLC35A4 and AltSLC35A4 do not play a role in cell fitness under normal conditions.

**Figure 5 :**
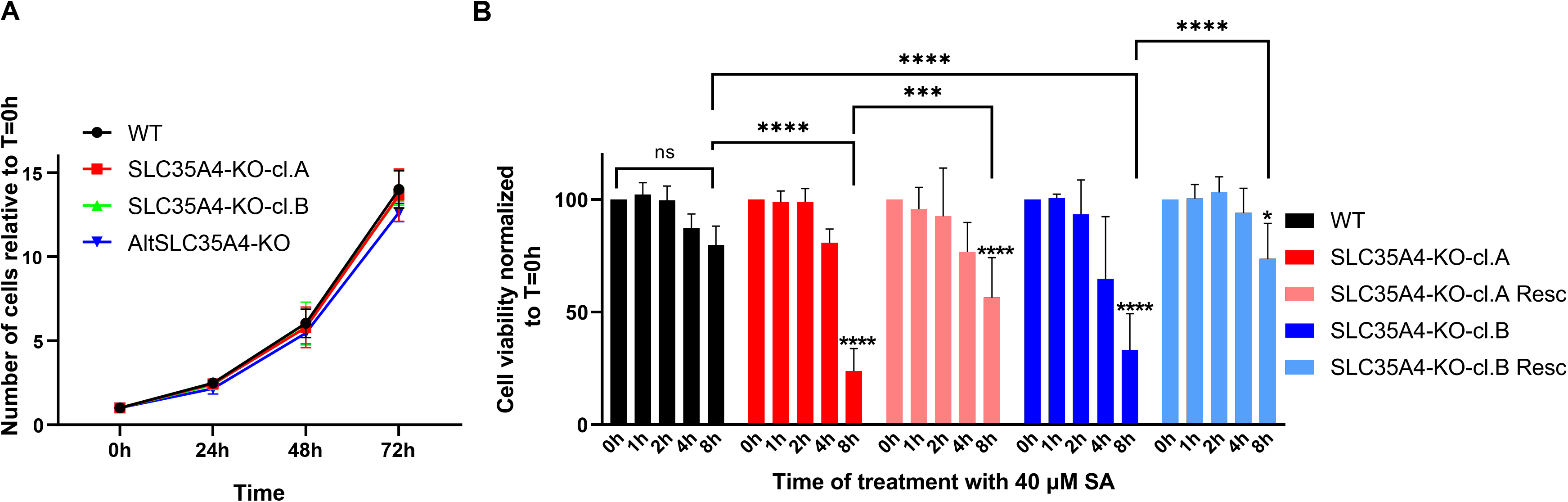
SLC35A4 is important for cell viability during cellular stress. **(A)** Cell proliferation curve of 143B cells (WT, SLC35A4-KO [clones A, B] and AltSLC35A4- KO). 5x10^4^ cells were seeded at T=0 h and counted daily over a period of 3 days. The results were normalized to the initial number of cells seeded. N=3 independent experiments. The significance of the results was assessed using a two-way ANOVA and Dunnett’s correction for multiple comparisons. **(B)** Cells of the indicated genotypes (WT, SLC35A4-KO-cl.A, cl.B, and their rescued counterparts stably expressing SLC35A4-3xHA) were treated with 40 µM sodium arsenite for the indicated times (0 h: vehicle). The ATP content of cells, reflecting cellular viability, was then quantified using the CellTiter-Glo 2.0 assay. The luminescence at 0 h for each cell genotype was defined as 100%. The significance of the results was determined using a two-way ANOVA and Tukey’s method for multiple comparisons. Asterisks placed directly above the graph’s bars indicate comparisons to T=0h within the same genotype. * P<0.05; ** P<0.01; **** P<0.0001. N=4 independent experiments.

Then, we tested whether the reference SLC35A4 is important for cellular resistance to OS by comparing cell viability of the KO lines compared to WT cells, in response to SA treatment. For that, we used the CellTiter-Glo 2.0 cell viability assay, that measures cellular ATP content to estimate viability. First, we examined the variations in cell viability within each genotype (Fig. 5B). WT cells showed a slight, but non-significant decrease in viability with increasing times of treatment, suggesting that the stress conditions were mild, allowing potential increased sensitivity to be revealed in other genotypes. In contrast, we observed a significant difference between the 0 h and 8 h time points in SLC35A4 KO cells, indicating that loss of SLC35A4 integrity sensitizes cells to stress.

When compared to WT cells, SLC35A4-KO cells (both cl. A and B) displayed a significant decrease in relative cell viability after 8 h of SA treatment. This effect was rescued by re-expressing SLC35A4-3xHA in SLC35A4-KO clones (Fig. 5B, Supplementary Fig. 1F). These results establish SLC35A4 as an important new player for cellular resistance to OS.

## DISCUSSION

Despite significant progress in genomics, current genome annotations continue to rely on arbitrary assumptions, such as the monocistronic nature of eukaryotic genes ^9^. Our study demonstrates that the *SLC35A4* is a bicistronic gene, encoding the reference protein SLC35A4 and the alternative protein AltSLC35A4. Our findings reinforce the growing understanding that alternative proteins, such as AltSLC35A4, are not merely incidental byproducts but hold significant functional potential. The biochemical assays employed, including subcellular fractionation, alkali treatment, and proteinase K protection assay, convincingly demonstrate AltSLC35A4’s specific insertion in the inner mitochondrial membrane. This confirmation clarifies previous conflicting reports by Yang et al., who identified its presence in the outer mitochondrial membrane (based on outer membrane localization of interactors of overexpressed Flag-tagged AltSLC35A4, and colocalization assay with various mitochondrial sub-compartments by confocal microscopy that has a limited resolution), and Rocha et al., who also located it in the inner mitochondrial membrane using biochemical assays on endogenous AltSLC35A4^21,29^. Localization of AltSLC35A4 in the inner mitochondrial membrane hints at a functional specialization that may contribute to mitochondrial homeostasis. Accordingly, Rocha et al. found that loss of AltSLC35A4 (a.k.a. SLC35A4-MicroProtein, SLC35A4-MP) leads to reduced basal, proton leak and maximal respiration in a rescuable manner, without altering expression of OXPHOS components. Conversely, overexpression of AltSLC35A4 modestly increased maximal respiratory capacity and mitochondrial membrane potential^21^. Future studies establishing the precise function of AltSLC35A4 within the mitochondrial inner membrane are needed.

To validate the results of Andreev et al. (2015) and Paolini et al. (2018) at the protein level, which indicate that SLC35A4 mRNA undergoes the highest increase in translational efficiency among all cellular mRNAs during OS and endoplasmic reticulum stress, we attempted to detect the endogenous protein. However, we encountered several challenges. Initially, we were hindered by the unavailability of a specific antibody. Indeed, only one publication^31^ reported using an antibody for SLC35A4; unfortunately, this antibody is no longer available. In addition, SLC solute carriers are notoriously difficult to study by WB, due to solubility and stability issues^32^. Furthermore, we were unable to detect SLC35A4 peptides using mass spectrometry, either at endogenous levels or following overexpression (SLC35A4-3xHA line, Fig. 3E, 4C), consistent with a recent report^21^.

To circumvent the difficulties in detecting the endogenous protein, we decided to generate stable cell lines that mimic the translation of the SLC35A4 protein from its endogenous mRNA.

To achieve this, we cloned the cDNA of the SLC35A4 gene, covering the entire sequence up to the stop codon of SLC35A4, thereby excluding the 3’ UTR region. This approach allows us to preserve the upstream open reading frame (uORF), which encompasses the sequence of the alternative protein (AltSLC35A4_SLC35A4-3xHA line). The rationale for this choice lies in the fact that, during cellular stress, the presence of a uORF has demonstrated regulatory potential for the reference protein^30,33,34^. Additionally, the translation of AltSLC35A4 is only slightly decreased during oxidative or endoplasmid reticulum stress^23,24^, consistent with our results showing no change in AltSLC35A4 levels (Fig. 3A,D).

In the context of the bicistronic construct (i.e. in a uORF-dependent manner), WB analysis revealed the appearance of SLC35A4 isoforms during OS, compatible with the increase in HA signal observed by immunofluorescence (Fig. 3A-C and Fig. 4A,B). This is likely due to ribosomal leaky scanning, a mechanism known to be facilitated by the presence of an uORF and the phosphorylation of eIF2α during cellular stress^23,30,35^. Our WB results were inconclusive regarding the dynamics of the full-length reference protein SLC35A4 expressed from the bicistronic construct, due to high variability in detected levels after OS induction. The increase in ribosome occupancy detected by Andreev et al. upon OS is observed across the whole SLC35A4 ORF^23^, not just more downstream regions that could be the source of shorter isoforms. Thus, leaky scanning could also facilitate full-length SLC35A4 expression to maintain SLC35A4 levels sufficient for adequate resilience to OS. Further investigation is needed to confirm the implication of leaky scanning, known to be modulated by the presence of uORFs. Notably, there were no changes induced by OS in cell lines lacking the uORF (monocistronic SLC35A4-3xHA construct), indicating that translational modulation of SLC35A4 (and its shorter isoforms) is dependent on the presence of the uORF and ruling out a role of proteolytic cleavage in short isoforms generation.

The integrated stress response (ISR) is a conserved cellular mechanism in eukaryotic cells, activated in response to internal or environmental stressors. It downregulates global protein synthesis while upregulating the expression of specific genes essential for the stress response^36^. The *SLC35A4* mRNA represents a clear example of this phenomenon, exhibiting the highest increase in translational efficiency among all cellular mRNAs during OS and endoplasmic reticulum stress^23,24^. In our study, we confirmed this increase at the protein level, and clarified this process by showing that appearance of shorter isoforms of SLC35A4 participates in the apparent translational increase. We also demonstrated that SLC35A4 is crucial for maintaining cell viability during OS, independently from the shorter isoforms since two SLC35A4-KO lines expressing the monocistronic SLC35A4-3xHA construct (that does not lead to shorter isoforms production in response to stress) were successfully rescued. The translational modulation of SLC35A4 necessitates the presence of an upstream open reading frame (uORF) during stress, similar to other ISR genes^37^. Whether the uORF-derived protein AltSLC35A4 also plays a role in cellular resistance to OS is currently under investigation. Nevertheless, mitochondria-localized AltSLC35A4 is unlikely to interact with its reference protein, localized in the Golgi under normal conditions.

We attribute the protective function of SLC35A4 to its role in transporting CDP-ribitol during cellular stress. Indeed, the transport of CDP-ribitol is primarily carried out by SLC35A1 under basal conditions^15^. In the context of the integrated stress response (ISR), the general translation of mRNAs, including SLC35A1, is inhibited^23^. Consequently, we hypothesize that SLC35A4, by escaping the translational downregulation of its mRNA, ensures the transport function of CDP-ribitol under cellular stress conditions. Moreover, ribitol has been shown to enhance nucleotide biosynthesis and increase levels of reduced glutathione^19^, which is involved in DNA synthesis and repair, the enhancement of vitamin C’s antioxidant activity, amino acid transport, and the detoxification of harmful compounds^38^. Additionally, CDP-ribitol supplementation has been associated with reduced muscle inflammation^39^, suggesting that CDP-ribitol possesses antioxidant and anti-inflammatory potential, thereby conferring cellular resistance to stress.

Furthermore, CDP-ribitol serves as a critical substrate for the glycosylation of specific proteins within the cell^39–41^. Studies have demonstrated that glycosylation affects OS by influencing the stability and function of antioxidant enzymes, modulating cell signaling pathways related to oxidative responses, and impacting protein folding and immune cell behavior, thereby playing a crucial role in cellular defense mechanisms against reactive oxygen species^42–45^. A plausible hypothesis is that OS could interfere with glycosylation in the event of CDP-ribitol deficiency; however, further studies are required to establish a direct link and prove this relationship.

In conclusion, we provide evidence for an important role of the SLC35A4 protein in protecting cells against oxidative stress, in a uORF-dependent manner. We confirm previous findings that AltSLC35A4 (or SLC35A4-MP), encoded from the 5’UTR of *SLC35A4*, is inserted in the mitochondrial inner membrane where it is likely to play important functions given its high degree of conservation in vertebrates. Future studies will unravel the precise function of this abundant alternative protein in mitochondrial homeostasis.

## ACKNOWLEDGEMENTS

We thank the members of the Vanderperre lab for fruitful discussions and technical help. Many thanks to the technical experts from CERMO-FC technological platforms (Denis Flipo and Grégoire Bonnamour - Cellular Analyses and Imaging; Geneviève Bourret – Genomics; Lekha Sleno and Léanne Ohlund – Metabolomics and Proteomics)

---

**Supplementary Figure 1:**

**(A)** PCR amplification of the CRISPR-targeted region for KO of AltSLC35A4, in WT cells and in the isolated KO clone (AltSLC35A4-KO). The appearance of a lower band in the KO clone indicates heterozygosity, which was confirmed by MiSeq sequencing and interpreted in (B).**(B)** Sequence alignment comparing the *in silico* translated alleles from the AltSLC35A4- KO clone to the WT protein sequence. **(C)** Immunofluorescence and confocal imaging showing AltSLC35A4-KO cells rescued with 3xHA-AltSLC35A4.The 3xHA-AltSLC35A4 was labeled with an anti-HA antibody (green). Nuclei were stained with Hoechst (blue). Scale bar: 100 µm. **(D)** PCR amplification of the CRISPR-targeted region for KO of SLC35A4, in WT cells and in the isolated KO clones A and B. Bands detected in the KO clones A and B indicate successful editing by deletion of a portion of the SLC35A4 coding sequence secondary to double cleavage at the target sequences of the two sgRNAs used. The specific allelic sequences were analyzed by Sanger sequencing and are interpreted in (E). **(E)** Sequence alignment comparing the *in silico* translated alleles of SLC35A4-KO clones A and B to the WT protein sequence. **(F)** Immunofluorescence and epifluorescence microscopy imaging validating the rescue of SLC35A4-KO clones A and B with the SLC35A4-3xHA monocistronic construct. The Golgi apparatus was labeled with an anti-Golgin97 antibody (green) and the SLC35A4-3xHA protein was labeled with an anti-HA antibody (magenta). Nuclei were stained with Hoechst (blue). Scale bars: 60 µm.

**Supplementary Figure 2:**

**(A)** Immunofluorescence and epifluorescence microscopy imaging of WT cells treated or not with 40 µM of SA for 3 hours. The stress marker ATF4 was labeled with an anti-ATF4 antibody (magenta). Nuclei were stained with Hoechst (blue). Scale bars: 50 µm. **(B)** Immunofluorescence and confocal imaging of AltSLC35A4_SLC35A4-3xHA expressing 143B cells labeled with an anti-HA antibody (green). Nuclei were stained with Hoechst (blue). Scale bars: 50 µm. **(C)** Immunofluorescence and epifluorescence microscopy imaging of SLC35A4-3xHA expressing 143B cells labeled with an anti-HA antibody (green). Golgi apparatus labeled with an anti-Golgin97 antibody (magenta) and nuclei were stained with Hoechst (blue). Scale bar : 30 µm.

**Supplementary Figure 3:**

**(A)** the substrate binding site (CDP-Ribitol) in SLC35A4 is shown. The circles in red represent key amino acids of the binding site deleted in both SLC35A4 KO clones. This diagram is modified from Ury et al., 2021^15^. (B) Illustrative diagram of WT (top) and the truncated (bottom) SLC35A4 protein produced from SLC35A4-KO clone B cells. TM: transmembrane domains; IH: interfacial alpha helix (IH) connecting TM 2 and 3. The consequences at the sequence level for SLC35A4-KO clone A cells are highly similar, where the mutated first and second alleles result in a 228 and 232 amino acids deletion, respectively (see Supplementary Figure 1D).

## Notes

### Competing Interest Statement

The authors have declared no competing interest.

## References

1. 1. International Human Genome Sequencing Consortium. Finishing the euchromatic sequence of the human genome. Nature 431, 931–945 (2004).

2. Cardon, T., Fournier, I. & Salzet, M. Shedding Light on the Ghost Proteome. Trends in Biochemical Sciences 46, 239–250 (2021).

3. Kapranov, P. et al. RNA Maps Reveal New RNA Classes and a Possible Function for Pervasive Transcription. Science 316, 1484–1488 (2007).

4. Wang, Z., Gerstein, M. & Snyder, M. RNA-Seq: a revolutionary tool for transcriptomics. Nat Rev Genet 10, 57–63 (2009).

5. Zhu, Y. et al. Discovery of coding regions in the human genome by integrated proteogenomics analysis workflow. Nat Commun 9, 903 (2018).

6. Vanderperre, B. et al. Direct Detection of Alternative Open Reading Frames Translation Products in Human Significantly Expands the Proteome. PLoS ONE 8, e70698 (2013).

7. Orr, M. W., Mao, Y., Storz, G. & Qian, S.-B. Alternative ORFs and small ORFs: shedding light on the dark proteome. Nucleic Acids Research 48, 1029–1042 (2020).

8. Wu, P. et al. Emerging role of tumor-related functional peptides encoded by lncRNA and circRNA. Mol Cancer 19, 22 (2020).

9. Brunet, M. A., Leblanc, S. & Roucou, X. Reconsidering proteomic diversity with functional investigation of small ORFs and alternative ORFs. Experimental Cell Research 393, 112057 (2020).

10. Mouilleron, H., Delcourt, V. & Roucou, X. Death of a dogma: eukaryotic mRNAs can code for more than one protein. Nucleic Acids Res 44, 14–23 (2016).

11. Vanderperre, B. et al. Direct Detection of Alternative Open Reading Frames Translation Products in Human Significantly Expands the Proteome. PLoS ONE 8, e70698 (2013).

12. Fredriksson, R., Nordström, K. J. V., Stephansson, O., Hägglund, M. G. A. & Schiöth, H. B. The solute carrier (SLC) complement of the human genome: phylogenetic classification reveals four major families. FEBS Lett 582, 3811–3816 (2008).

13. Hadley, B. et al. Nucleotide Sugar Transporter SLC35 Family Structure and Function. Comput Struct Biotechnol J 17, 1123–1134 (2019).

14. Eckhardt, M., Gotza, B. & Gerardy-Schahn, R. Mutants of the CMP-sialic acid transporter causing the Lec2 phenotype. J Biol Chem 273, 20189–20195 (1998).

15. Ury, B., Potelle, S., Caligiore, F., Whorton, M. R. & Bommer, G. T. The promiscuous binding pocket of SLC35A1 ensures redundant transport of CDP-ribitol to the Golgi. Journal of Biological Chemistry 296, 100789 (2021).

16. Brändli, A. W., Hansson, G. C., Rodriguez-Boulan, E. & Simons, K. A polarized epithelial cell mutant deficient in translocation of UDP-galactose into the Golgi complex. Journal of Biological Chemistry 263, 16283–16290 (1988).

17. Guillen, E., Abeijon, C. & Hirschberg, C. B. Mammalian Golgi apparatus UDP-N-acetylglucosamine transporter: molecular cloning by phenotypic correction of a yeast mutant. Proc Natl Acad Sci U S A 95, 7888–7892 (1998).

18. Ortiz-Cordero, C. et al. NAD+ enhances ribitol and ribose rescue of α-dystroglycan functional glycosylation in human FKRP-mutant myotubes. eLife 10, e65443 (2021).

19. Tucker, J. D., Doddapaneni, R., Lu, P. J. & Lu, Q. L. Ribitol alters multiple metabolic pathways of central carbon metabolism with enhanced glycolysis: A metabolomics and transcriptomics profiling of breast cancer. PLOS ONE 17, e0278711 (2022).

20. Ibraghimov-Beskrovnaya, O. et al. Primary structure of dystrophin-associated glycoproteins linking dystrophin to the extracellular matrix. Nature 355, 696–702 (1992).

21. Rocha, A. L. et al. An inner mitochondrial membrane microprotein from the SLC35A4 upstream ORF regulates cellular metabolism. Journal of Molecular Biology 168559 (2024) doi:10.1016/j.jmb.2024.168559.

22. Fulda, S., Gorman, A. M., Hori, O. & Samali, A. Cellular Stress Responses: Cell Survival and Cell Death. International Journal of Cell Biology 2010, e214074 (2010).

23. Andreev, D. E. et al. Translation of 5′ leaders is pervasive in genes resistant to eIF2 repression. eLife 4, e03971 (2015).

24. Paolini, N. A. et al. Ribosome profiling uncovers selective mRNA translation associated with eIF2 phosphorylation in erythroid progenitors. PLoS ONE 13, e0193790 (2018).

25. Lu, P. D., Harding, H. P. & Ron, D. Translation reinitiation at alternative open reading frames regulates gene expression in an integrated stress response. The Journal of Cell Biology 167, 27–33 (2004).

26. Jourdain, A. A. et al. GRSF1 Regulates RNA Processing in Mitochondrial RNA Granules. Cell Metabolism 17, 399–410 (2013).

27. Pagliarini, D. J. et al. A Mitochondrial Protein Compendium Elucidates Complex I Disease Biology. Cell 134, 112–123 (2008).

28. Vanderperre, B. et al. MPC1-like Is a Placental Mammal-specific Mitochondrial Pyruvate Carrier Subunit Expressed in Postmeiotic Male Germ Cells. Journal of Biological Chemistry 291, 16448–16461 (2016).

29. Yang, H., Li, Q., Stroup, E. K., Wang, S. & Ji, Z. Widespread stable noncanonical peptides identified by integrated analyses of ribosome profiling and ORF features. Nat Commun 15, 1932 (2024).

30. Andreev, D. E., et al. TASEP modelling provides a parsimonious explanation for the ability of a single uORF to derepress translation during the integrated stress response. Elife 7, e32563 (2018).

31. Sosicka, P. et al. An insight into the orphan nucleotide sugar transporter SLC35A4. Biochimica et Biophysica Acta (BBA) - Molecular Cell Research 1864, 825–838 (2017).

32. Tsuji, Y. Transmembrane protein western blotting: Impact of sample preparation on detection of SLC11A2 (DMT1) and SLC40A1 (ferroportin). PLoS ONE 15, e0235563 (2020).

33. Moro, S. G., Hermans, C., Ruiz-Orera, J. & Albà, M. M. Impact of uORFs in mediating regulation of translation in stress conditions. BMC Mol Cell Biol 22, 29 (2021).

34. Andreev, D. E., Dmitriev, S. E., Zinovkin, R., Terenin, I. M. & Shatsky, I. N. The 5’ untranslated region of Apaf-1 mRNA directs translation under apoptosis conditions via a 5’ end-dependent scanning mechanism. FEBS Lett 586, 4139–4143 (2012).

35. Baird, T. D. & Wek, R. C. Eukaryotic Initiation Factor 2 Phosphorylation and Translational Control in Metabolism12. Adv Nutr 3, 307–321 (2012).

36. Pakos-Zebrucka, K., et al. The integrated stress response. EMBO reports 17, 1374–1395 (2016).

37. Andreev, D. E. et al. Translation of 5′ leaders is pervasive in genes resistant to eIF2 repression. eLife 4, e03971 (2015).

38. Montecinos, V. et al. Vitamin C Is an Essential Antioxidant That Enhances Survival of Oxidatively Stressed Human Vascular Endothelial Cells in the Presence of a Vast Molar Excess of Glutathione*. Journal of Biological Chemistry 282, 15506–15515 (2007).

39. Wu, B. et al. Ribitol dose-dependently enhances matriglycan expression and improves muscle function with prolonged life span in limb girdle muscular dystrophy 2I mouse model. PLoS One 17, e0278482 (2022).

40. Kanagawa, M. & Toda, T. Ribitol-phosphate—a newly identified posttranslational glycosylation unit in mammals: structure, modification enzymes and relationship to human diseases. The Journal of Biochemistry 163, 359–369 (2018).

41. Cataldi, M. P., Lu, P., Blaeser, A. & Lu, Q. L. Ribitol restores functionally glycosylated α-dystroglycan and improves muscle function in dystrophic FKRP-mutant mice. Nat Commun 9, 3448 (2018).

42. An, Y. et al. The role of oxidative stress in diabetes mellitus-induced vascular endothelial dysfunction. Cardiovascular Diabetology 22, 237 (2023).

43. Medina-Navarro, R., Durán-Reyes, G., Díaz-Flores, M. & Vilar-Rojas, C. Protein Antioxidant Response to the Stress and the Relationship between Molecular Structure and Antioxidant Function. PLoS One 5, e8971 (2010).

44. Gerlach, J. Q., Sharma, S., Leister, K. J. & Joshi, L. A Tight-Knit Group: Protein Glycosylation, Endoplasmic Reticulum Stress and the Unfolded Protein Response. in Endoplasmic Reticulum Stress in Health and Disease (eds. Agostinis, P. & Afshin, S.) 23–39 (Springer Netherlands, Dordrecht, 2012). doi:10.1007/978-94-007-4351-9_2.

45. Li, S. et al. Role of O-linked N-acetylglucosamine protein modification in oxidative stress-induced autophagy: a novel target for bone remodeling. Cell Communication and Signaling 22, 358 (2024).

