## Supplementary figures and images for "The dual coding gene *SLC35A4* protects against oxidative stress"

### Supplementary figures

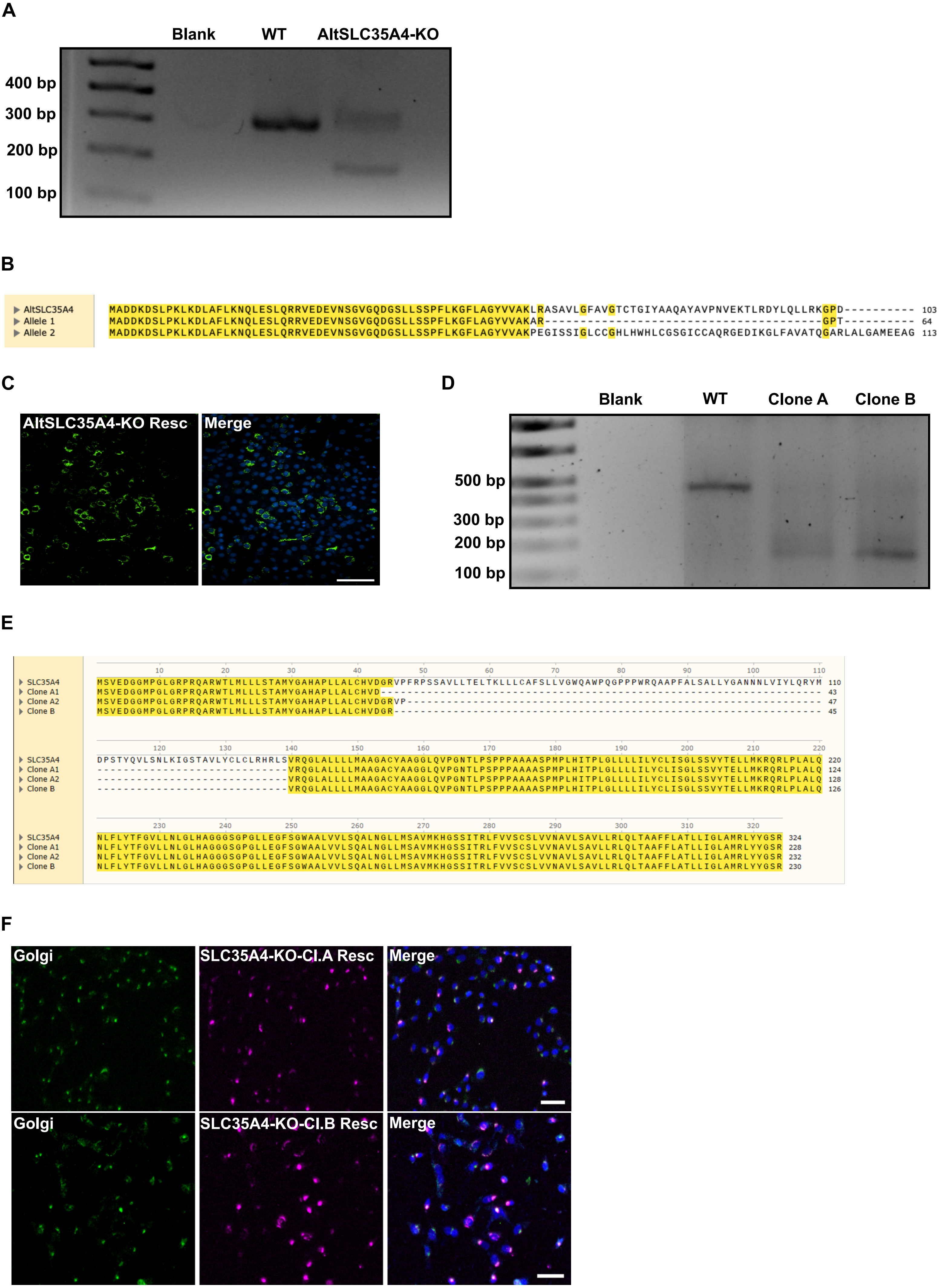

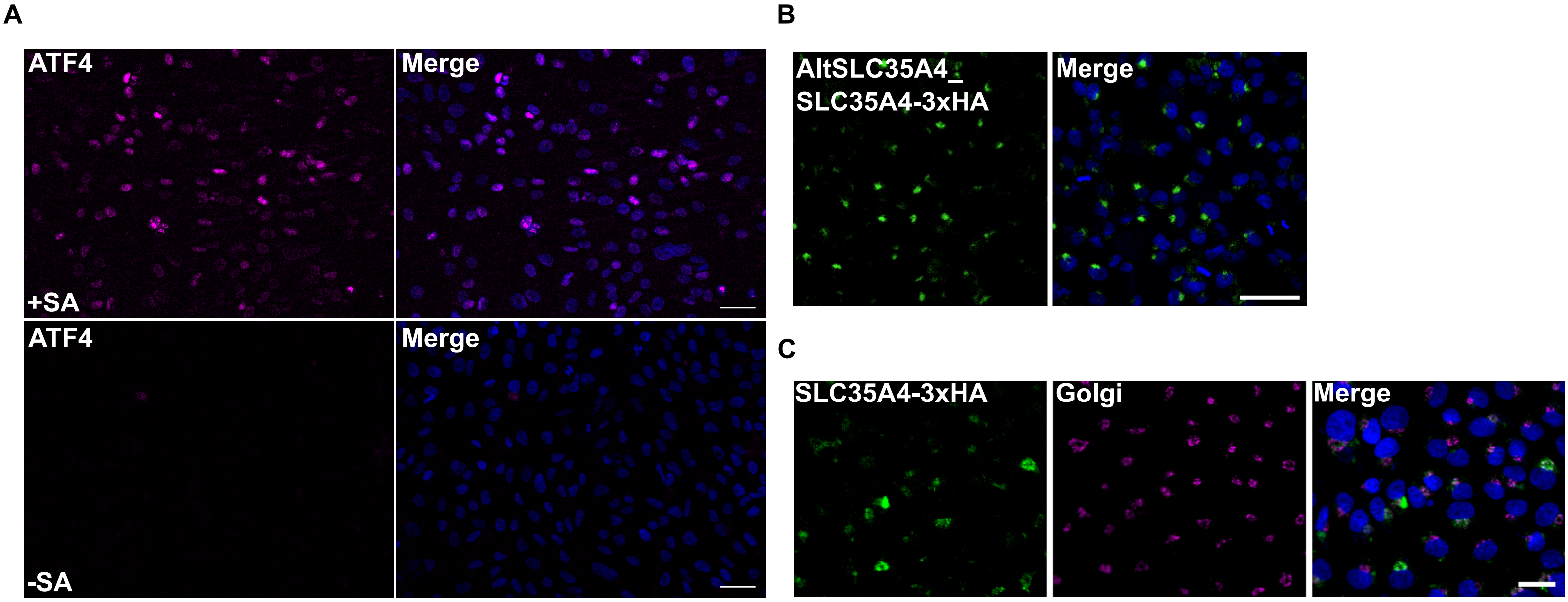

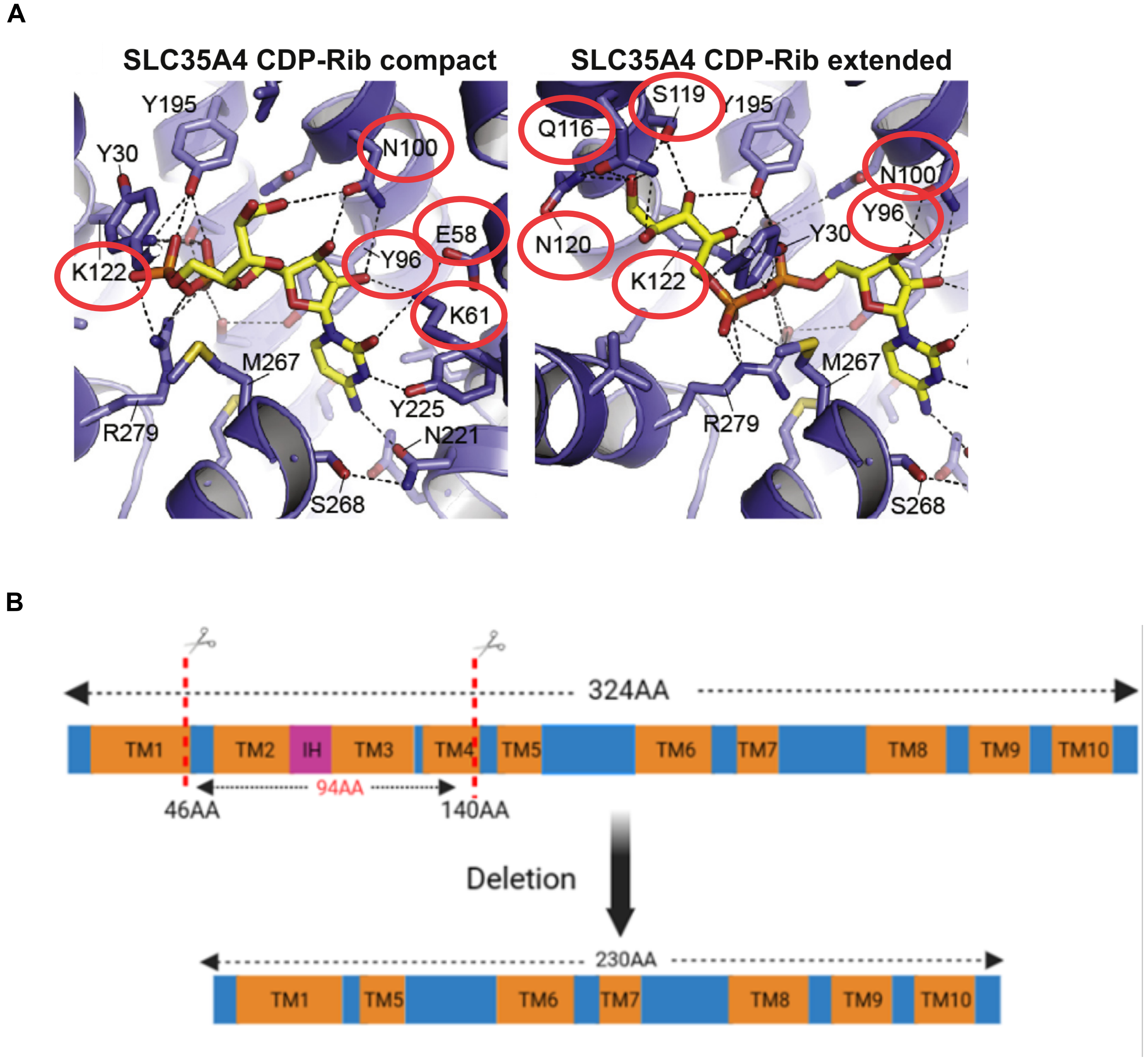
